# Self-organizing actin networks drive sequential endocytic protein recruitment and vesicle release on synthetic lipid bilayers

**DOI:** 10.1101/2023.02.14.528546

**Authors:** Emily H. Stoops, Michael A. Ferrin, Danielle M. Jorgens, David G. Drubin

## Abstract

Forces generated by actin assembly assist membrane invagination during clathrin-mediated endocytosis (CME). The sequential recruitment of core endocytic proteins and regulatory proteins, and assembly of the actin network, are well documented in live cells and are highly conserved from yeasts to humans. However, understanding of CME protein self-organization, as well as the biochemical and mechanical principles that underlie actin’s role in CME, is lacking. Here, we show that supported lipid bilayers coated with purified yeast WASP, an endocytic actin assembly regulator, and incubated in cytoplasmic yeast extracts, recruit downstream endocytic proteins and assemble actin tails. Time-lapse imaging of WASP-coated bilayers revealed sequential recruitment of proteins from different endocytic modules, faithfully replicating *in vivo* behavior. Reconstituted actin networks assemble in a WASP-dependent manner and deform lipid bilayers, as seen by electron microscopy. Time-lapse imaging revealed that vesicles are released from the lipid bilayers with a burst of actin assembly. Actin networks pushing on membranes have previously been reconstituted; here, we have reconstituted a biologically important variation of these actin networks that self-organize on bilayers and produce pulling forces sufficient to bud off membrane vesicles. We propose that actin-driven vesicle generation may represent an ancient evolutionary precursor to diverse vesicle forming processes adapted for a wide array of cellular environments and applications.

**Significance Statement:** Actin filament assembly participates in many vesicle-forming processes. However, the underlying principles for how assembly is initiated and organized to effectively harness assembly forces remain elusive. To address this gap, we report a novel reconstitution of actin-driven vesicle release from supported lipid bilayers. Using real-time imaging, we observe sequential recruitment of endocytic proteins and, following a burst of actin assembly, vesicle release from bilayers. Given the absence of cargo or upstream endocytic regulatory proteins on the bilayers, and the participation of actin in many vesicle-forming processes, we posit that this mode of vesicle formation represents an early evolutionary precursor for multiple trafficking pathways. We expect that this assay will be of great use for future investigations of actin-mediated vesicle-forming processes.

## Introduction

Clathrin-mediated endocytosis (CME) involves the recruitment of dozens of endocytic proteins to the plasma membrane where they collect cargo, deform the membrane, and drive vesicle scission. Endocytic protein recruitment follows a precise order and timing (1, 2), and can be roughly subdivided into a variable early phase during which clathrin and adapter components arrive at the membrane, and a highly regular late phase during which Wiskott Aldrich Syndrome Protein (WASP) and actin-associated proteins act at the endocytic site (3). Recruitment of *S. cerevisiae* WASP (Las17, referred to hereafter as scWASP) to a threshold level promotes a burst of actin assembly through activation of the Arp2/3 complex, which nucleates actin polymerization (4). scWASP-mediated actin assembly is required to overcome the high turgor pressure of the yeast cell and drive membrane invagination (5).

With over 60 proteins involved in yeast endocytosis, many with non-essential roles or additional non-endocytic cellular functions, determining the functions of each protein *in vivo* has been challenging (6, 7). CME robustness often thwarts attempts at function elucidation; deletion of seven genes encoding early endocytic proteins was shown to only minimally impact endocytosis (8). *In vivo* studies are well suited to identify necessary components of the CME machinery, but to identify components that are sufficient for CME and to gain insights into their biochemical mechanisms, we sought to reconstitute endocytic events *in vitro*. Biochemical reconstitution of cellular processes provides otherwise unattainable insight into fundamental molecular mechanisms (9). Reconstitution assays have been irreplaceable to test models and reveal mechanistic details for processes from pathogen motility in infected host cells (10–12) to mitotic spindle formation (13, 14). Of relevance to the work reported here, many important advances in our understanding of vesicle fusion and fission have come through *in vitro* reconstitutions (15–18). A reconstituted system for actin-mediated CME promises to reveal principles for properly ordered recruitment of CME proteins and the mechanics of membrane vesicle formation.

Here, we developed an *in vitro* assay that faithfully replicates the cooperative assembly of endocytic proteins onto synthetic membranes and deformation of the membrane into a nascent vesicle, resulting in actin-driven vesicle formation and release. We demonstrate that scWASP-templated actin and endocytic protein network assembly alone are sufficient to direct membrane deformation and vesiculation from lipid bilayers. We suggest that WASP-mediated actin network assembly represents a minimal system to drive membrane invagination and vesicle scission and may represent an early evolutionary precursor for a wide variety of membrane trafficking modalities.

## Results and Discussion

An *in vitro* actin assembly assay using polystyrene microbeads functionalized with scWASP was developed previously (19). When incubated in yeast cell extract, scWASP-coated beads assembled actin and selectively recruited endocytic actin network proteins. A fraction of the beads became motile. While this earlier reconstitution study represented an important advance for understanding the cooperative assembly of actin and actin-binding proteins involved in endocytosis, it did not allow conclusions to be drawn about the effect of the membrane on assembly of the actin network, or of actin network assembly on the membrane. To examine the effects of endocytic protein assembly on membranes we modified this system to include lipid bilayers (Figure 1A). To that end, supported lipid bilayers were made on solid microspheres using established protocols that retain bilayer fluidity (Figure S1A). scWASP, previously shown to activate the Arp2/3 complex *in vitro* (20, 21), was purified and efficiently recruited to supported lipid bilayers (Figure 1B, 1C).

**Figure 1.**
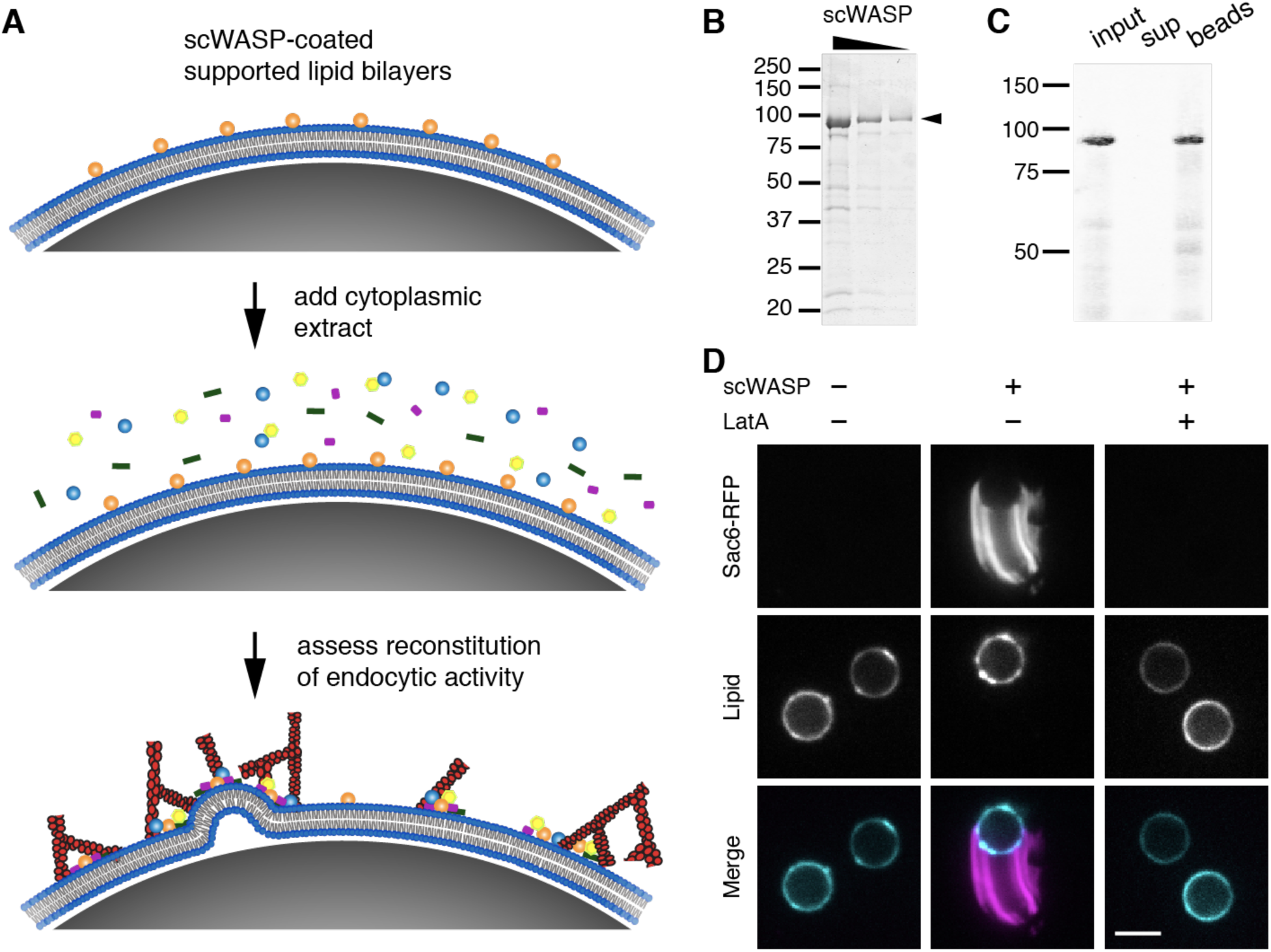
Reconstitution of endocytic actin networks on supported lipid bilayers. (A) Schematic representation of the *in vitro* reconstitution assay. Purified scWASP (Las17) is attached to supported lipid bilayers, which are then incubated in yeast cytoplasmic extract, resulting in the assembly of endocytic actin networks. (B) GelCode-stained SDS-PAGE gel of purified scWASP (arrowhead). (C) Purified scWASP binds to supported lipid bilayers, as shown by Western blot probed with an anti-Las17 antibody. (D) Sac6-RFP from yeast cytoplasmic extract labels actin networks assembled on scWASP-coated supported lipid bilayers (Atto647-DOPE, lipid). 180 µM latrunculin A (LatA) inhibits actin assembly on bilayers. Scale bar, 5 µm.

Cytoplasmic extracts were prepared from yeast strains expressing at endogenous levels endocytic proteins tagged with fluorescent proteins (Table S1). scWASP-coated bilayers were incubated in freshly generated cytoplasmic extract and were then imaged.

### Reconstitution of endocytic actin networks on lipid bilayers in yeast extract

As previously observed with microbeads lacking a membrane bilayer, scWASP-coated supported lipid bilayers accumulated dense actin networks and associated actin binding proteins (Figure 1D). Membrane-associated filamentous actin was detected using GFP-tagged Sac6 (yeast fimbrin, an actin-binding protein, Figure 1D) and fluorescent phalloidin, a small molecule that binds to filamentous actin (Figure S1B). Approximately 30% of beads broke symmetry and became motile, forming an actin tail behind the lipid-coated bead. 3D reconstructions show tube-like actin structures trailing lipid-coated motile beads, with little Sac6 signal inside the tube (Movie S1). These structures resemble those previously observed in bacterial motility reconstitution in extracts (11). The actin networks associated with non-motile beads failed to break symmetry but continued to assemble actin over time (Movie S2). Actin assembly on bilayers was dependent on the presence of scWASP (Figure 1D). Addition of the actin monomer sequestering drug latrunculin A (LatA) to reactions prohibited actin tail assembly and bead motility.

Previous work demonstrated roles for anionic lipid species in CME site initiation and invagination, including phosphatidylserine (PS) and phosphatidylinositol 4,5 bisphosphate (PI(4, 5)P_2_) (22). Varying the lipid composition to include PS and/or PI(4, 5)P_2_ at concentrations used in previous bilayer reconstitution systems (17, 23–25) did not result in a pronounced effect on actin assembly on the beads, as measured by quantifying the percentage of beads that accumulated a Sac6 signal (Figure S1C and S1D).

### scWASP-coated bilayers assemble the machinery that carries out CME internalization

To assess how faithfully the reconstitution system recapitulated *in vivo* CME events, we determined whether proteins downstream of scWASP in the endocytic pathway were recruited to the bilayers (Figure 2A). We used extracts from yeast strains expressing a GFP-tagged protein of interest and Sac6-RFP to allow endocytic proteins to be located relative to the actin network.

**Figure 2.**
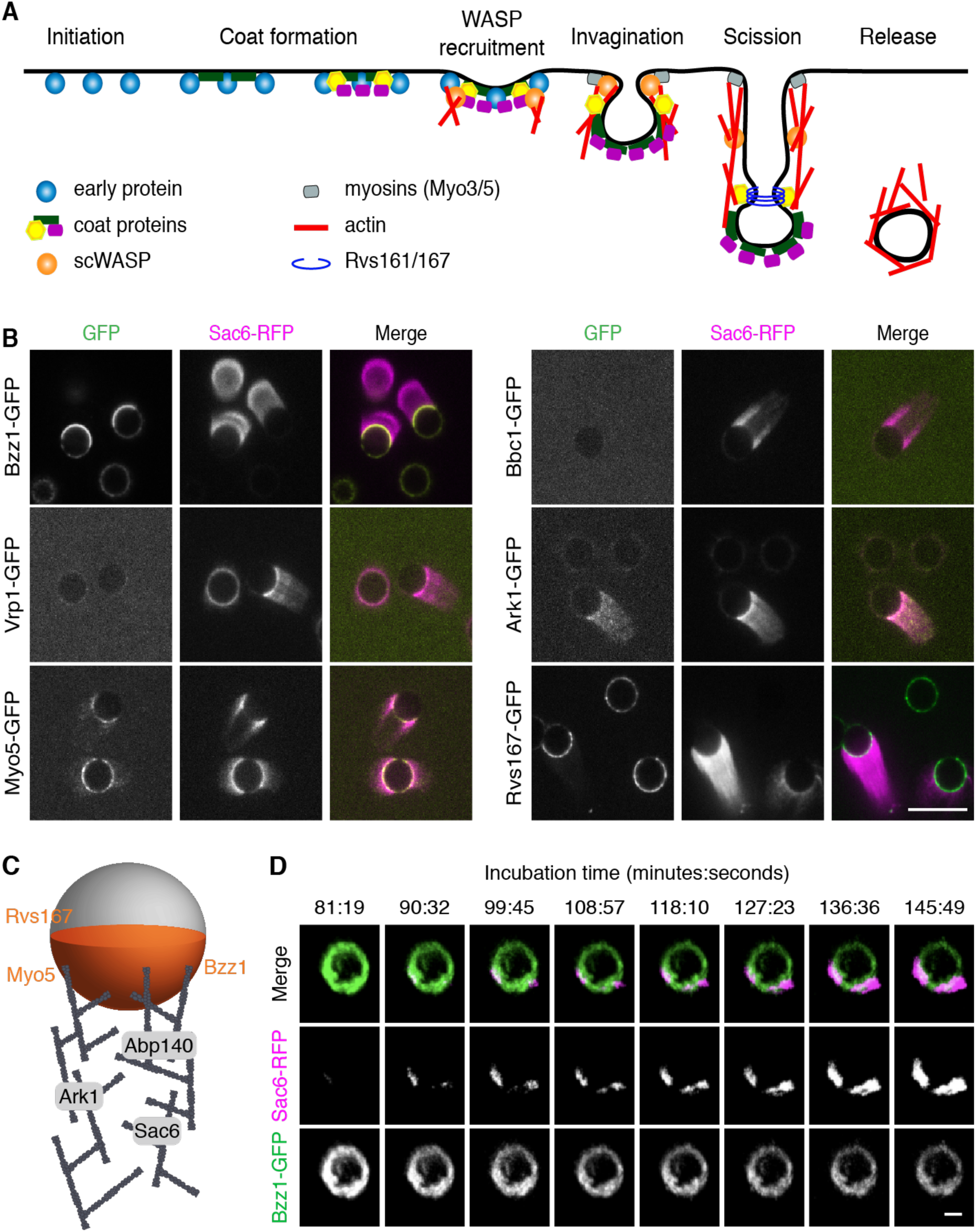
scWASP-coated bilayers sequentially recruit downstream endocytic proteins. (A) Schematic representation of clathrin-mediated endocytosis in *S. cerevisiae*. (B) Protein localization on scWASP-coated PC/PS supported bilayers. Extracts were prepared from strains expressing Sac6-RFP and the indicated endocytic proteins tagged with GFP. Representative fluorescence images from 3 independent experiments are shown. Scale bar, 10 µm. (C) Localization of endocytic proteins on supported lipid bilayers and associated actin networks is represented in cartoon view. Bzz1, Rvs167, and Myo5 were observed primarily on lipid bilayers in the region of actin assembly. Ark1, Sac6, and Abp140 were associated with the actin network. (D) scWASP-coated beads were incubated in cytoplasmic extract from strains expressing Bzz1-GFP and Sac6-RFP. Representative montage of 3D rendered images from time-lapse confocal microscopy. Indicated time points (minutes:seconds) are relative to the moment cytoplasmic extract was added to beads. Scale bar, 2 µm.

Myo3 and Myo5 are type I myosins that facilitate endocytic actin assembly through their motor and NPF (nucleation promoting factor) activities (26–28). They are required for endocytic vesicle internalization (21, 26, 29, 30). Myo5-GFP was localized primarily to the bilayers in areas of actin assembly, with trace amounts of Myo5-GFP also localizing to actin tails (Figure 2B). These data are consistent with previous results on scWASP-coated microbeads (19) and with previous data showing that type I myosin does not leave the cell surface during vesicle internalization (31). Bzz1 was also recruited to bilayers in an scWASP-dependent manner (Figures 2B and S2A). Bzz1 is an scWASP-binding protein that relieves scWASP inhibition to promote actin assembly initiation (21, 32). On lipid bilayers with actin tails, Bzz1-GFP localized to the hemisphere of the lipid bilayer from which the actin tail protrudes. In contrast, Vrp1 (Wasp Interacting Protein [WIP]), which interacts directly with scWASP and Myo3/5 (21), was not observed or was only weakly observed on bilayers or associated with actin tails. Similarly, Bbc1 was not recruited to bilayers or actin tails under these conditions. Bbc1 is known to inhibit scWASP in later stages of endocytosis and to thereby control the speed and extent of inward vesicle movement (21, 30). Bbc1 absence from actin tails might reflect incomplete reconstitution of regulatory components. Following scWASP activation there is a burst of actin assembly and recruitment of actin-associated proteins. As observed for Sac6-GFP, actin-binding protein Abp140-GFP was recruited to actin tails (Figure S2B). Additionally, the endocytic regulatory serine/threonine protein kinase Ark1 was observed in actin tails protruding from scWASP-coated lipid bilayers (Figures 2B and S2B).

Further downstream of scWASP, Rvs161/Rvs167 heterodimeric N-BAR protein (yeast homologues of amphiphysin) contributes to vesicle scission and has been shown to tubulate membranes *in vitro* (30, 33). Observations in electron micrographs show that Rvs161/Rvs167 is concentrated at the bud neck of deeply invaginated clathrin-coated pits (34, 35). In our assay, Rvs167-GFP was recruited to scWASP-coated bilayers and was concentrated in the area of actin network formation, where it formed punctae (Figure 2B).

We also determined whether proteins upstream of scWASP are recruited to the membranes. Sla2 links coat proteins to the actin cytoskeleton in vivo. However, it was not visible on scWASP-coated bilayers (Figure S2C).

These data are summarized in a cartoon schematic showing where each endocytic protein localizes on the scWASP-coated lipid bilayers (Figure 2C). Our results are like those obtained previously with non-lipid-coated beads (19). The recruitment of multiple actin binding proteins, nucleation promoting factors, and actin regulatory components to the bilayers or actin tails, or both, indicates that our reconstituted system recapitulates self-assembly of the endocytic actin network. Furthermore, Rvs167 presence indicates recruitment of at least some of the machinery required to induce vesicle scission.

### Protein recruitment order *in vitro* recapitulates CME temporal dynamics *in vivo*

As a test of whether the correct CME protein recruitment order is encoded in the proteins themselves, we determined whether this sequence is recapitulated in the reconstituted system. The *in vivo* recruitment of proteins to endocytic sites follows a regular order that is highly conserved across species (1, 3, 36, 37) (Figure 2A). To determine whether CME proteins accumulate on lipid bilayers in the correct physiological sequence, we observed reconstituted endocytic protein complexes as they formed in a flow chamber over time. Figure 2D shows time-lapse data from scWASP-coated beads after addition of cytoplasmic extract prepared from cells expressing Bzz1-GFP and Sac6-RFP. In cells, Bzz1 is present at the endocytic site immediately prior to the start of actin assembly (21, 32). Our *in vitro* data recapitulate this sequence with Bzz1 appearing on beads before Sac6 (Figure 2D, Movie S2). Additionally, Bzz1 recruitment is scWASP-dependent (Figure S2B), indicating that the established sequence from scWASP to the regulatory factor (Bzz1) to actin assembly is preserved. Interestingly, Bzz1 localization on the bead is polarized prior to actin assembly, which then initiates from areas of high Bzz1 density.

These data demonstrate successful recapitulation of sequential endocytic protein assembly and establish that the spatiotemporal recruitment of endocytic proteins is an intrinsic property of the proteins. It should be noted that while the recruitment order is preserved *in vitro*, the kinetics of recruitment do not replicate *in vivo* kinetics. In live cells, Bzz1 and Sac6 have lifetimes of approximately 15-17 and 11-15 seconds at the endocytic site, respectively (3, 21, 30). On supported lipid bilayers, Bzz1 is recruited to bilayers within seconds of extract addition, but there is a delay of up to 90 minutes prior to Sac6 arrival. The variance in length of time between recruitment events in the cytosolic extract versus the native yeast cytosolic environment could reflect differences in the protein concentrations and energy conditions, or differences between length scales for *in vivo* endocytic events, which are on the nanometer scale, versus *in vitro* length scales for actin tails on the order of microns.

### Vesicles are released from sites of endocytic actin network assembly

The supported lipid bilayer system described above reconstitutes actin-generated pushing forces on the micron scale, as demonstrated by bead motility. We next set out to determine whether these components could also drive nano-scale vesicle formation. In experiments similar to those described above to test for sequential CME protein recruitment, we next used a small amount of Atto647-DOPE as a marker of the lipid bilayer. The micrographs revealed tubules and smaller lipid protuberances in regions where endocytic proteins and actin assembled on the bilayers (Figure S3A, S3B). In addition to tubulation, we often observed in static confocal micrographs small (diffraction-limited) lipid structures embedded in, or in the vicinity of, actin tails (Movies S3 and S4), which we hypothesized to be vesicles released from the supported-lipid bilayer. To determine whether these putative vesicles form by being pinched off from supported lipid bilayers, real-time imaging was performed. We systematically tested the effect of PS and PI(4, 5)P_2_ in lipid bilayers on vesicle accumulation in our assay. Addition of 5% PI(4, 5)P_2_ to our standard lipid mixture produced the highest frequency of actin-associated vesicles (Table S2). In 37 time-lapse videos collected from four independent experiments, we observed spherical lipid structures breaking free from the lipid bilayer concurrent with a burst of actin polymerization at the same site (Figure 3A, Movie S5). The number of vesicles observed per bead (0.248 ± 0.291) was reduced dramatically by removing either scWASP (0.0476 ± 0.0638) or by treating reactions with LatA (0.0481 ± 0.0638); no vesicles were observed in the absence of cytoplasmic extract (Table S3). To quantitatively analyze the dynamics of these vesiculation events and compare them to *in vivo* CME, we tracked vesicle formation and movement in 3D and measured fluorescence intensities and displacement over time (Figures 3B and S4B). Each vesicle track was temporally aligned at the displacement inflection point calculated using previously described techniques (Figure S5, (38)). This alignment revealed a highly reproducible burst of actin assembly beginning approximately 15 minutes before the vesicle buds off from the bead (Figure 4C). The actin network continues to grow and propel the vesicle from the bead about 2 µm on average over about 10 minutes (Figure 4C). This time scale is significantly longer than is seen in yeast, where the time from onset of actin assembly to vesicle scission is approximately 30 seconds. It is possible that ultracentrifugation and dilution of the cytoplasmic extract alter its composition sufficiently to affect reaction kinetics. Alternatively, the assay may be reconstituting friction-mediated scission (39) in a system with less friction than in the native plasma membrane. Despite the difference in time scale, the sequence from scWASP to Bzz1 recruitment to actin assembly (as visualized by Sac6) to membrane pulling follows the order observed for endocytic events *in vivo*.

**Figure 3.**
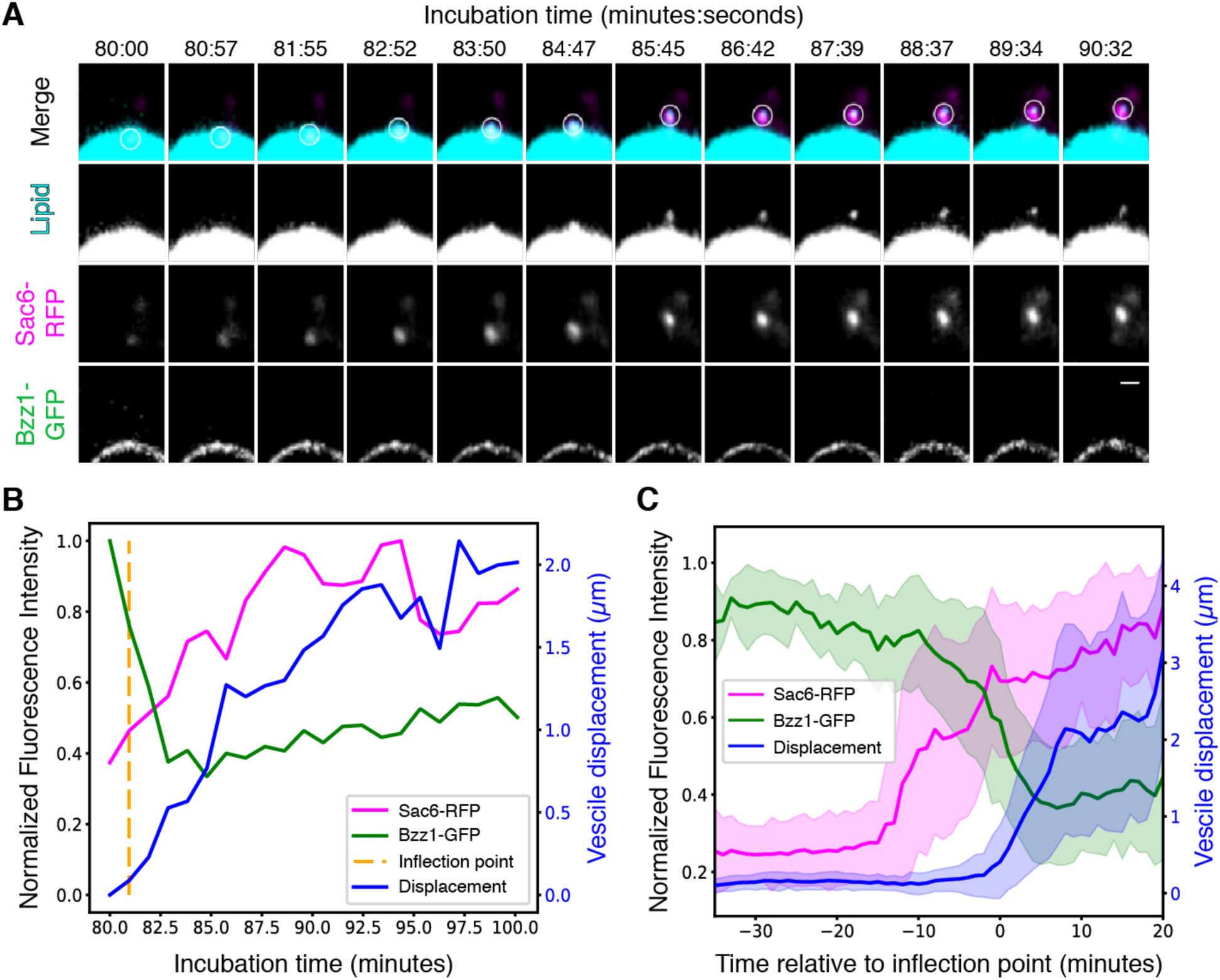
Reconstitution of actin-mediated vesicle budding. Supported lipid bilayers were coated with scWASP and incubated in Bzz1-GFP Sac6-RFP cytoplasmic extracts for the indicated times. (A) Montage of a single vesiculation event. A maximum intensity projection 2 µm in depth is displayed for all channels. Bilayers include 2% Atto647-DOPE (cyan) to allow visualization of lipid. The white circle in the merged channel row indicates the region used for quantitative analysis. Scale bar, 1 µm. (B) Quantification trace of fluorescence intensities and vesicle displacement from the vesiculation event visualized in (A). The algorithmically determined inflection point is indicated by the vertical dotted line. (C) Mean fluorescence intensities and vesicle displacement with standard deviation from all measured vesiculation events (n=37). All traces were aligned in time by their respective inflection points prior to taking the mean and standard deviation.

**Figure 4.**
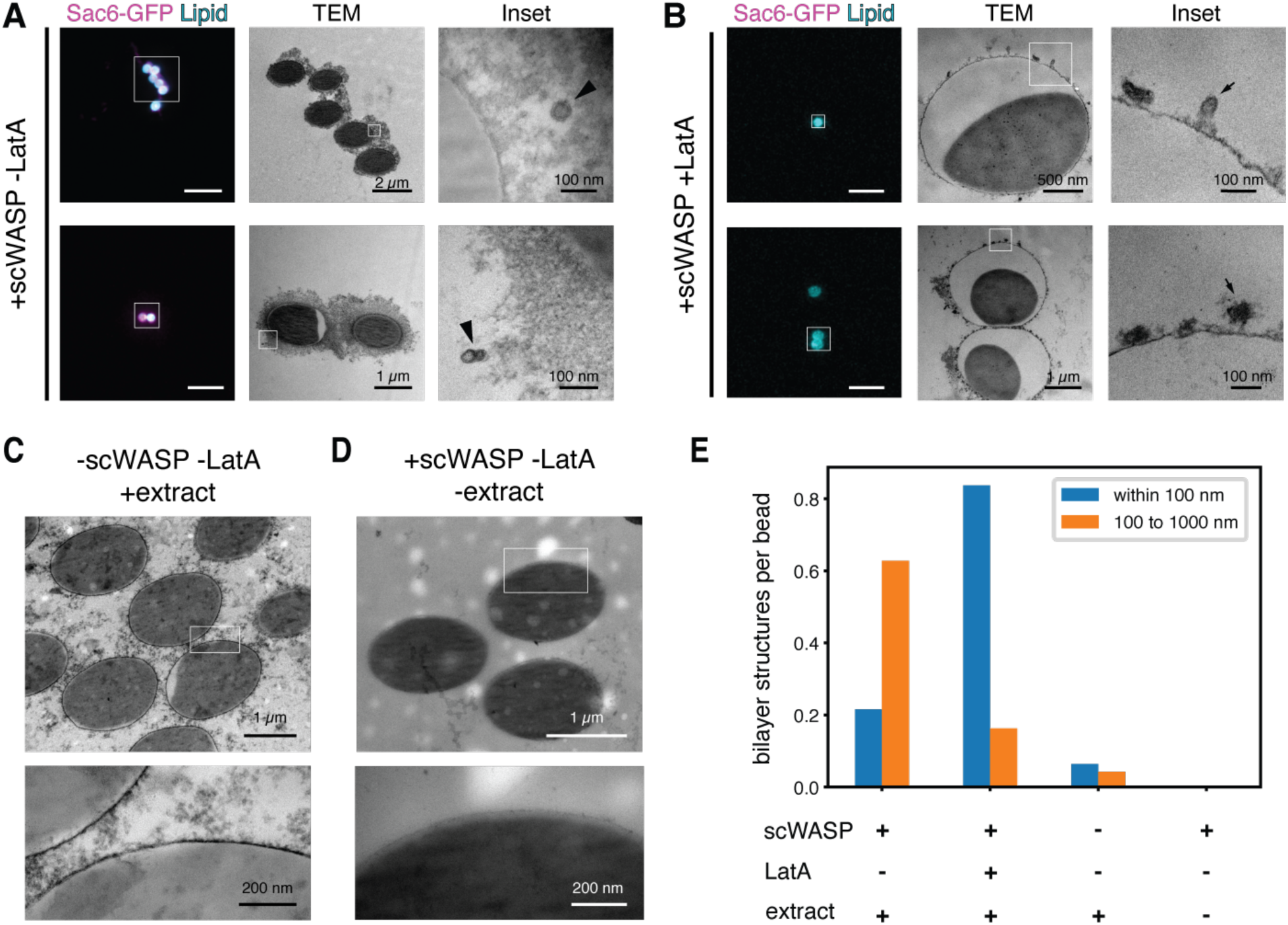
Self-assembled endocytic actin networks deform lipid bilayers. (A and B) Representative CLEM images of scWASP-coated supported lipid bilayers incubated in extracts containing Sac6-GFP in the (A) presence or (B) absence of latrunculin A. Left subpanels show light microscopy (scale bar, 10 µm). Middle and right subpanels show transmission EM images of the regions indicated by white boxes. (C and D) Representative electron micrographs of supported bilayer reactions in the absence of (C) scWASP or (D) cytoplasmic extract. (E) Manual scoring of lipid bilayer tubules and vesicles in EM sections prepared from the indicated reaction conditions.

Given that endocytic events typically occur on a length scale that is below the diffraction limit of our fluorescence microscopy system, we employed correlative light and electron microscopy (CLEM) to achieve the necessary resolution to directly observe deformed membranes and free vesicles. We used beads 2 μm in diameter (versus 5 μm in previous experiments) to improve the beads’ spatial stability during sample processing for CLEM. Vesiculation events occurred on both size beads at similar rates (Figure S4, Movies S6 through S8). Transmission electron micrographs of lipid bilayers on beads in yeast extract revealed formation of scWASP-dependent membrane structures on the length scale of biological endocytic events (∼50-100nm, Figures 4A and S6). A variety of membrane profiles were observed, including apparent tubules contiguous with the bead surface and vesicles some distance away from the bead (Figures 4A and S6). Corresponding light micrograph images demonstrated actin assembly around the beads (Figure 4A, Sac6-GFP signal). To assess whether membrane deformation requires actin assembly, we sampled bilayers treated with latrunculin A and observed a pronounced reduction in free vesicles, but an increase in small tubules and deformations at the membrane surface (Figure 4B). Fewer free vesicles and membrane deformations were observed in samples lacking scWASP (Figure 4C) and these features were entirely absent in samples lacking cytoplasmic extract (Figure 4D). We lack 3D information in these TEM micrographs to allow us to determine whether the membrane structures are contiguous with the supported lipid bilayer. However, we have observed that vesicles move away from the beads in time-lapse imaging. Therefore, as a proxy for whether they are contiguous with the membrane or are distinct structures, we classified vesicles observed in EM by their distance from the bead surface (Figure 3E, Table S2). We observed an approximately 15-fold increase in the frequency of membrane structures between 100 and 1000 nm from the surface of scWASP-coated beads compared to the surface of beads lacking scWASP. Membranes observed in this region are likely vesicles derived from the supported lipid bilayer on the bead surface. Similar to our quantification of the time-lapse imaging data (Table S3), more vesicles were observed in reactions containing scWASP and extract in the absence of latrunculin A. Of note, scWASP-containing beads treated with latrunculin A to inhibit actin polymerization showed the highest frequency of membrane structures within 100 nm of the bead surface. These structures are likely invaginations of the bead supported lipid bilayer. Interestingly, the curvature-generating protein Bzz1 is recruited to supported lipid bilayers in a scWASP-dependent manner, independent of latrunculin A treatment (Figure S6C). These data are consistent with a role for curvature generating proteins in early stages of membrane invagination and a known role for actin in driving membrane internalization and scission in yeast.

## Conclusion

This work demonstrates that endocytic proteins and the actin network intrinsically embody the information sufficient to self-organize into a force-producing system capable of deforming a lipid bilayer and producing a vesicle. This work builds off other *in vitro* models for actin force generation, including those for *Listeria* motility (10), filopodia formation (40, 41) and actin-induced membrane phase separation (42). However, unlike previously reported systems, here actin is generating a pulling, not pushing, force. Moreover, time-lapse imaging revealed that the order of molecular events was faithfully reconstituted, and electron microscopy showed that these membrane remodeling events are on the size scale observed in yeast endocytosis.

Interestingly, vesicle generation in our reconstituted system occurs in the apparent absence of coat proteins such as clathrin, and in the absence of certain upstream structural and regulatory factors including Sla2 and Bbc1. Our data are consistent with previous findings, which showed that the coat is not necessary for endocytic vesicle formation. Endocytosis still occurs in clathrin null mutants, although the number of endocytic sites is reduced (30, 43–45). A previous study examined CME in yeast with null alleles of genes encoding seven initiation factors and coat proteins and determined that membrane bending and vesicle budding were largely unperturbed (8). These observations are consistent with the existence of multiple endocytic pathways in various cell types that require actin but do not require clathrin or its associated coat proteins (46, 47). In light of these observations, we propose that a network of actin and associated proteins might constitute a primitive endocytic machinery that diversified through evolution to give rise to multiple vesicle-forming processes, with coat proteins and other accessory factors added later as evolutionary embellishments. We anticipate that coat proteins would be required to reconstitute selective cargo recruitment. In the future, addition of cargo and proteins upstream of scWASP in the CME pathway to this *in vitro* reconstitution assay promises to provide valuable mechanistic insights into the roles of cargo, coat proteins and other regulatory factors.

This work establishes that vesicle pulling, and scission, are intrinsic to the ensemble activities of the endocytic actin network. Future work using this assay will allow us to elucidate the roles of individual proteins and lipid species in vesicle generation.

## Materials and Methods

### Strains

Cells were maintained at 25 or 30°C on rich YPD media (<u>Y</u>east extract/<u>P</u>eptone/<u>D</u>extrose). Yeast strains used in this study are listed in Table S1. Genomic C-terminal tagging was performed as described previously (48).

### Protein purification

Yeast WASP (Las17) was cloned into the p363 overexpression plasmid using PacI and XmaI restriction sites to be in frame with a C-terminal TEV site, 3x StreptagII, and 9x His tag. The recombinant protein was overexpressed in yeast using a GAL induction system in the D1074 strain and purified as described previously (49). Briefly, protein expression was induced with 2% galactose for 6 to 8 hrs at 30°C. Following induction, cells were flash frozen. Frozen cells were lysed by mechanical shearing in a cryogenic grinder (SPEX SamplePrep). The resulting powder was reconstituted in buffer containing protease inhibitors (Roche) and centrifuged at 80k rpm for 20 min at 4°C. The supernatant was passed through a 0.45 µm syringe filter and then applied to a HisTrap column. Samples containing Las17 were pooled and dialyzed in a Slide-a-Lyzer cassette (Thermo Fisher) after addition of TEV protease. After a brief incubation with Ni-agarose to remove the protease and cleaved His tag, Las17 was concentrated in an Ultra-4 centrifugal filter device (Amicon). Aliquots of purified Las17 were flash frozen in liquid N_2_.

### Supported lipid bilayer production

Supported lipid bilayers were prepared using a previously published protocol with minor modifications (50). Briefly, chloroform stocks of lipids (Avanti Polar Lipids) were combined in a glass vial etched with Piranha (3:1 H_2_SO_4_/H_2_O_2_) according to the molar ratios listed for each experiment. The lipid mixture was dried at 42°C under vacuum using a rotary evaporator. After brief exposure to nitrogen, the dried lipid film was rehydrated in deionized water to 1 mg/mL. The mixture was passed through a microextruder (Avanti Polar Lipids) fitted with a 0.1μm filter 11 times to generate small unilamellar vesicles (SUVs). SUVs were used immediately or stored for up to 5 days at 4°C.

Polystyrene or silica microspheres (Bangs Laboratories) were cleaned for 15 minutes in 1% Helmenex, followed by three wash steps in ddH_2_O. SUVs diluted 1:5 in 10x TBS (to a final concentration of 2x TBS) were incubated with the microbeads, generating supported lipid bilayers through self-assembly. Unruptured vesicles were removed via multiple wash steps in TBS.

### Generation of whole cell lysates

Cells were grown in YPD at 25°C to an OD_600_ of 0.6-0.8, as measured on an Ultrospec 10 Cell Density Meter (Amersham). Cells from 4 L of culture were harvested by centrifugation, washed in cold water, and centrifuged again. Standing moisture was removed from pellets and cells were flash frozen in liquid N_2_ and then stored at -80°C. Frozen cultures were lysed by mechanical shearing in a pre-chilled cryogenic grinder (SPEX SamplePrep) using a medium-sized SPEX vial that had been pre-chilled in liquid nitrogen. Milling consisted of a 5 min pre-chill followed by 6-10 cycles comprising 3 min of grinding and 1 min of rest. The sample vial remained submerged in liquid N_2_ throughout the milling process. Powdered lysate was collected in a 50 ml conical vial that had been pre-chilled in liquid N_2_. Lysate preparations stored at -80°C were stable for >6 months but were highly sensitive to temperature excursions.

### Yeast extract preparation

HK buffer (40mM HEPES, pH 7.5, 200mM KCl, 1mM PMSF) was supplemented with cOmplete mini EDTA-free protease inhibitor cocktail (Roche), according to manufacturer specifications. Powdered lysate was weighed out into a pre-chilled 5 mL glass beaker using a pre-chilled spatula. 875μl of supplemented HK buffer at 4°C was added to each gram of yeast powder. After thawing on ice, samples were centrifuged in a pre-chilled polycarbonate ultracentrifuge tube for 25 min at 345,000 x g at 4°C. The cleared supernatant was collected immediately following the centrifuge run using a syringe to transfer it to a pre-chilled 1.5 ml tube and was then used in an assay within 10 minutes of the final ultracentrifugation step. Care was taken to avoid disturbing either the pellet or the white lipid layer. Using extract longer than 10 minutes following centrifugation or generating extract from powdered lysate that had not been consistently kept either at -80°C or in liquid N_2_ were both found to negatively impact reconstitution.

### Membrane functionalization and bead assay

180 nM yeast WASP (Las17) was incubated on the membranes for 1 h. Beads were washed and then incubated in 1% casein for 15 min to block nonspecific binding. After washing to remove excess casein, beads were stored in 1x TBS for up to 1 hour. 1-2 μl of functionalized bilayers were added to cytosolic extract to a total reaction volume of 20 μl. 1 mM ATP and, where noted, 180 µM latrunculin A (Sigma Aldrich) were added immediately to the bilayer and extract mixture. Microbeads were spotted on slides and imaged within 1 hour.

### Microscopy and image analysis

#### Fluorescence microscopy

Except where noted otherwise, all images were acquired on Nikon Eclipse Ti inverted Yokogawa spinning disk confocal microscope fitted with Andor CSU-X spinning disc confocal equipment and controlled by Nikon Elements software. Imaging was performed using a 100x 1.45 NA Plan Apo λ oil immersion objective and an Andor IXon X3 EM-CCD camera. GFP, RFP, and Atto647 fluorescence was excited using 488-, 561- and 638-nm lasers, respectively. Images for figure panel S1C were collected using a Nikon Eclipse Ti microscope equipped with a 100x 1.4 NA Plan Apo VC oil objective and an Andor Neo sCMOS camera. The system was controlled using Metamorph software (Molecular Devices). Images for figure panels 4A (lower subpanel) and 4B (both subpanels) were acquired on a AxioObserver Zeiss LSM 710 Laser Scanning Confocal with 40x 1.4 NA Plan Apo oil immersion objective and PMT detector. GFP and Atto647 fluorescence was excited using 488- and 633-nm lasers, respectively. The system was controlled with Zen 2010 software. All imaging devices were kept in rooms maintained at 23 to 25°C.

#### Fluorescence recovery after photobleaching

Membrane fluidity was assessed using fluorescence recovery after photobleaching (FRAP) performed on a Zeiss LSM 710 confocal microscope fitted with a Plan-Apochromat 63x 1.40 NA oil immersion DIC M27 and Zen software. An ROI drawn over a portion of the bilayer was bleached using 594 nm laser light at 100% laser power for 50 iterations. Images were taken every 780 ms before and after bleaching. Images were analyzed using ImageJ software. The plot profile function in Fiji was used to measure the signal intensity of TexasRed-DHPE in the bleached ROI and the opposite (unbleached) side of the bilayer. FRAP data were normalized to the highest and lowest mean values in the pre-bleach condition.

#### Correlative light and electron microscopy

Supported lipid bilayers were generated on polystyrene microbeads and incubated with 500nM scWASP and cytosolic extract, as described above. Reactions were fixed in 4% paraformaldehyde, 0.05% glutaraldehyde for at least 30 min at 4°C. Beads were deposited on poly-L-lysine coated 35 mm gridded glass-bottom dishes (MatTek). After washing, adherent beads were embedded in 2% low melting point agarose (Electron Microscopy Sciences) and submerged in 1x TBS for microscopy. Regions of interest were identified by Atto647-DOPE and Sac6-GFP fluorescence signal, and then mapped to spatial coordinates on the gridded coverslip by brightfield imaging. Samples were then washed in 1x PBS and stained with 1% osmium tetroxide and 1.6% potassium ferricyanide at 4°C for 45 min. Following additional washes with PBS, and a quick exchange in water, samples were dehydrated with an ascending gradient of ethanol followed by pure ethanol before they were progressively infiltrated with resin and left overnight in unaccelerated Epon resin (Ted Pella). Epon resin with accelerant was exchanged onto the sample three times, then the dishes were incubated at 60°C for 16-20 hours for resin polymerization. Following polymerization, the glass coverslips were removed using ultra-thin Personna razor blades (EMS, Hatfield, PA, USA). Correlative light and electron microscopy was performed to visualize specific beads within regions of interest. Regions of interest, identified by the gridded alpha-numerical labeling on the plates were carefully removed, precisely trimmed to the area of interest, and mounted on a blank resin block with cyanoacrylate glue for sectioning. Serial thin sections (80 nm) were cut using a Leica UC6 ultramicrotome (Leica, Wetzlar, Germany) from the surface of the block until approximately 4-5 microns in to ensure complete capture of the bead. Section-ribbons were then collected sequentially onto formvar-coated slot or 50 mesh grids. The grids were post-stained with 2% uranyl acetate followed by Reynold’s lead citrate, for 5 min each. The sections were imaged using a FEI Tecnai 12 120kV TEM (FEI, Hillsboro, OR, USA) and data recorded using either a Gatan US1000 CCD with Digital Micrograph 3 or a Gatan Rio 16 CMOS with Gatan Microscopy Suite software (Gatan Inc., Pleasanton, CA, USA). Images were adjusted for brightness and contrast in ImageJ (National Institutes of Health), and rotated, but otherwise unaltered.

#### Image analysis

Images were processed using ImageJ software. Pixel intensity values were rescaled identically for all images from an experiment. To allow visualization of small structures, some pixels in some images were intentionally saturated. To quantify Sac6 accumulation on lipid bilayers of varying compositions, beads were manually counted in the DIC channel and then scored for presence or absence of Sac6 fluorescence.

Additional image processing of time-lapse spinning disc confocal fluorescence microscopy data was performed using ImageJ plugins following recommendations by Picco et al (51). First, background subtraction was applied to raw data from each channel using a rolling ball algorithm with 100 pixel radius. We then performed bleach correction with an exponential decay model on each individual channel except when imaging Sac6-RFP, because for this marker there was no decay in fluorescence signal over the course of our time-lapses. Individual beads were then cropped and processed with a 3D drift correction algorithm so that positions of vesicles could later be measured relative to their source bead. The resulting multi-channel volumetric time-lapse processed data were then rendered in 3D and assembled into montages using custom python scripts (available on GitHub [https://github.com/DrubinBarnes/Stoops_Ferrin_et_al_2023]).

To quantify vesiculation dynamics, we used the TrackMate plugin of ImageJ to manually track the position of vesicles in time-lapse data after the processing steps described in the previous paragraph. We measured the 3D displacement of each tracked vesicle from its starting position over time, as well as the average fluorescence intensity of a 0.4 µm radius sphere around the center of each vesicle over time. To collectively analyze data for all tracked vesicles, we first aligned each individual trajectory in time by the inflection point of the displacement curve (adapted from (38)). The inflection point was defined as the time point at which the vesicle displacement was the maximum negative difference from a linear regression fit to each vesiculation event trace, out of all the time points before the point at the maximum positive difference. Aligned fluorescence intensity curves were normalized to the maximum value for each vesicle before calculating the average and standard deviation for all tracked vesicles.

## Acknowledgments

We thank Michelle Lu, Ross Pedersen, and members of the Drubin/Barnes laboratory for frequent informal discussions. We thank Akemi Kunibe and Damien D’Amours for generously providing strains and plasmids. We thank Holly Aaron and Feather Ives for microscopy training and assistance. Thank you to Reena Zalpuri and the staff at the University of California Berkeley Electron Microscope Laboratory for assistance in electron microscopy sample preparation and data collection. Confocal microscopy was conducted at the University of California, Berkeley, Cancer Research Laboratory Molecular Imaging Center, supported by the Gordon and Betty Moore Foundation. This work was supported by National Institutes of Health grants R35GM118149 to D. G. Drubin and 1F32GM113383-01A1 to E. H. Stoops, as well as the National Science Foundation Graduate Research Fellowship Program to M. A. Ferrin. Research reported in this publication was supported in part by the National Institutes of Health S10 program under award number 1S10RR026866-01. The content is solely the responsibility of the authors and does not necessarily represent the official views of the National Institutes of Health. The authors declare no competing financial interests.

---

**Movie S1. 3D rendering of actin tail assembled on scWASP-coated bilayer.** Related to Figure 1D. scWASP-coated supported lipid bilayers were incubated in cytoplasmic extract expressing Sac6-RFP. A 3D rendering of a confocal stack is displayed. Blue outline indicates bead location. Scale bars, 5 µm.

**Movie S2. Time-lapse and 3D rendering of sequential recruitment of endocytic proteins to scWASP-coated supported bilayers.** Related to Figure 2D. scWASP-coated supported lipid bilayers were incubated in Bzz1-GFP (green) Sac6-RFP (magenta) cytoplasmic extracts. A 3D rendering of a substack of the bead is displayed. Time after addition of extract to bilayers is displayed. Frames were generated every 57 s and are played back at 6.7 fps. Scale bars, 5 µm.

**Movie S3. 3D rendering of a vesicle embedded in an actin tail.** Related to Figure S3. scWASP-coated supported lipid bilayers were incubated in cytoplasmic extracts from cells expressing Bzz1-GFP (not displayed) and Sac6-RFP (magenta). Bilayers include 2% Atto647-DOPE (cyan) to allow visualization of lipid. A 3D rendering of a static confocal stack of the bead is displayed. The white circle indicates location of a diffraction-limited vesicle embedded in an actin tail. Scale bars, 5 µm.

**Movie S4. 3D rendering of an actin-associated vesicle in the vicinity of a supported lipid bilayer.** Related to Figure S3. scWASP-coated supported lipid bilayers were incubated in cytoplasmic extracts from cells expressing Bzz1-GFP (not displayed) and Sac6-RFP (magenta). Bilayers include 2% Atto647-DOPE (cyan) to allow visualization of lipid. A 3D rendering of a static confocal stack of the bead is displayed. The white circle indicates the location of diffraction-limited, actin-associated vesicles. Scale bars, 5 µm.

**Movie S5. 3D rendering of reconstitution of actin-mediated vesicle budding.** Related to Figure 3A. scWASP-coated supported lipid bilayers were incubated in Bzz1-GFP (green) Sac6-RFP (magenta) cytoplasmic extracts. Bilayers include 2% Atto647-DOPE (cyan) to allow visualization of lipid. A 3D rendering of a substack of the bead is displayed. The white circle indicates the region of vesicle budding. A white line traces the path of the vesicle. Time after addition of extract to bilayers is displayed. Frames were generated every 57 s and are played back at 6.7 fps. Scale bars, 5 µm.

**Movie S6. 3D rendering of actin-mediated vesicle budding from supported lipid bilayer on 5 µm bead.** Related to Figure S4, upper panel. scWASP-coated supported lipid bilayers were incubated in Bzz1-GFP (green) Sac6-RFP (magenta) cytoplasmic extracts. Bilayers include 2% Atto647-DOPE (cyan) to allow visualization of lipid. A 3D rendering of a substack of the bead is displayed. The white circle indicates the region of vesicle budding. A white line traces the path of the vesicle. Time after addition of extract to bilayers is displayed. Frames were generated every 30.8 s and are played back at 6.7 fps. Scale bars, 5 µm.

**Movie S7. 3D rendering of actin-mediated vesicle budding from supported lipid bilayer on 5 µm bead.** Related to Figure S4, middle panel. scWASP-coated supported lipid bilayers were incubated in Bzz1-GFP (green) Sac6-RFP (magenta) cytoplasmic extracts. Bilayers include 2% Atto647-DOPE (cyan) to allow visualization of lipid. A 3D rendering of a substack of the bead is displayed. The white circle indicates the region of vesicle budding. A white line traces the path of the vesicle. Time after addition of extract to bilayers is displayed. Frames were generated every 79 s and are played back at 6.7 fps. Scale bars, 5 µm.

**Movie S8. 3D rendering of actin-mediated vesicle budding from supported lipid bilayer on 2 µm bead.** Related to Figure S4, lower panel. scWASP-coated supported lipid bilayers were incubated in Bzz1-GFP (green) Sac6-RFP (magenta) cytoplasmic extracts. Bilayers include 2% Atto647-DOPE (cyan) to allow visualization of lipid. A 3D rendering of a substack of the bead is displayed. The white circle indicates the region of vesicle budding. A white line traces the path of the vesicle. Time after addition of extract to bilayers is displayed. Frames were generated every 62.8 s and are played back at 6.7 fps. Scale bars, 5 µm.

**Figure S1.**
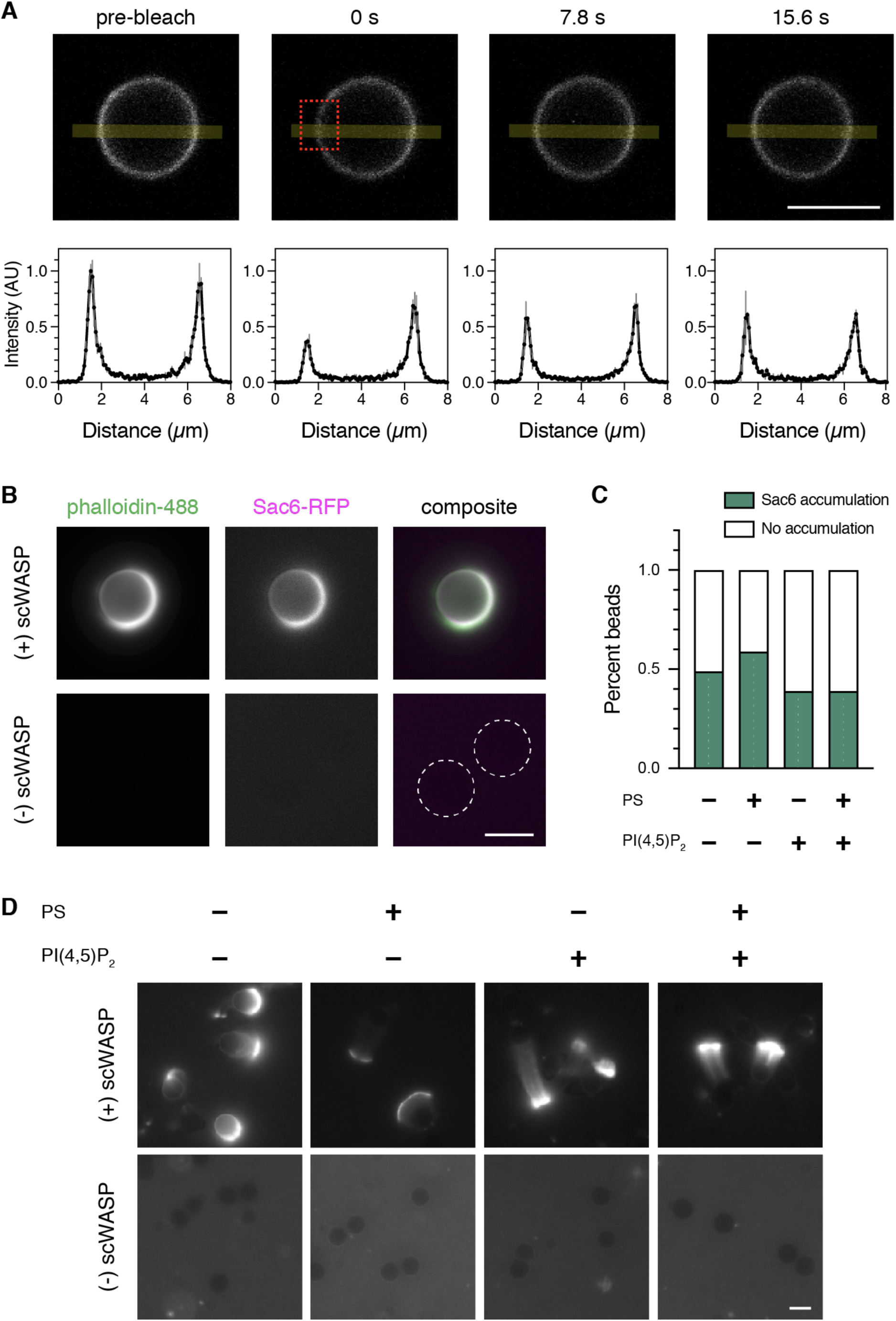
(A) Fluorescence recovery after photobleaching experiment of supported lipid bilayers containing 0.05% TexasRed-DHPE. Time-lapse confocal images and the corresponding fluorescence intensity of a line (yellow line) drawn through the bleaching region (red box) demonstrate rapid lipid turnover. (B) Phalloidin-488 (26.4µM) was added to scWASP-coated or uncoated bilayers (75% PC, 20% PS, 5% DGS-NTA) during incubation with cytoplasmic extract containing Sac6-RFP. Dotted lines outline the location of the beads lacking scWASP. (C and D) Sac6-GFP from yeast cytoplasmic extract labels actin networks assembled on scWASP-coated supported bilayers. Bilayer composition included PS and/or PI(4,5)P_2_ as indicated. (C) Quantification of the percentage of beads with associated Sac6 fluorescence. At least 160 beads per sample were quantified. (D) Representative images of Sac6-GFP accumulation on bilayers. Scale bars, 5 µm.

**Figure S2.**
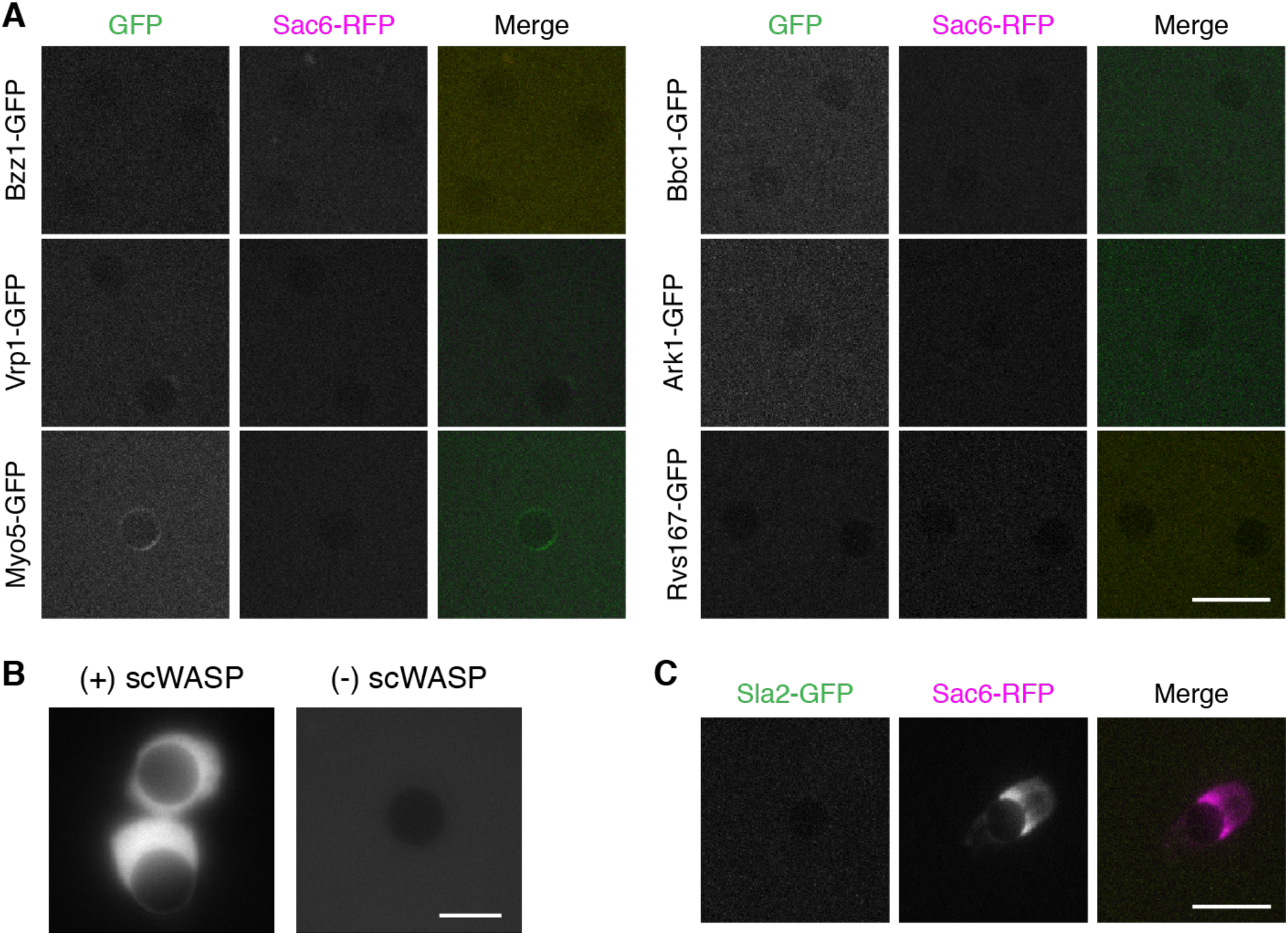
(A) Endocytic proteins do not localize to bilayers in the absence of scWASP. Extracts were prepared from strains expressing Sac6-RFP and the indicated endocytic proteins tagged with GFP, as in Figure 2B. Scale bar, 10 µm. (B) Abp140-GFP is recruited to actin networks. scWASP-coated and uncoated bilayers were incubated in cytoplasmic extract generated from a strain expressing Abp140-GFP. Scale bar, 5 µm. (C) Coat protein Sla2 does not localize to scWASP-coated bilayers. Bilayers were incubated in cytoplasmic extract generated from a strain expressing Sla2-GFP and Sac6-RFP. Scale bar, 10 µm. Representative fluorescence images from 3 independent experiments are shown.

**Figure S3.**
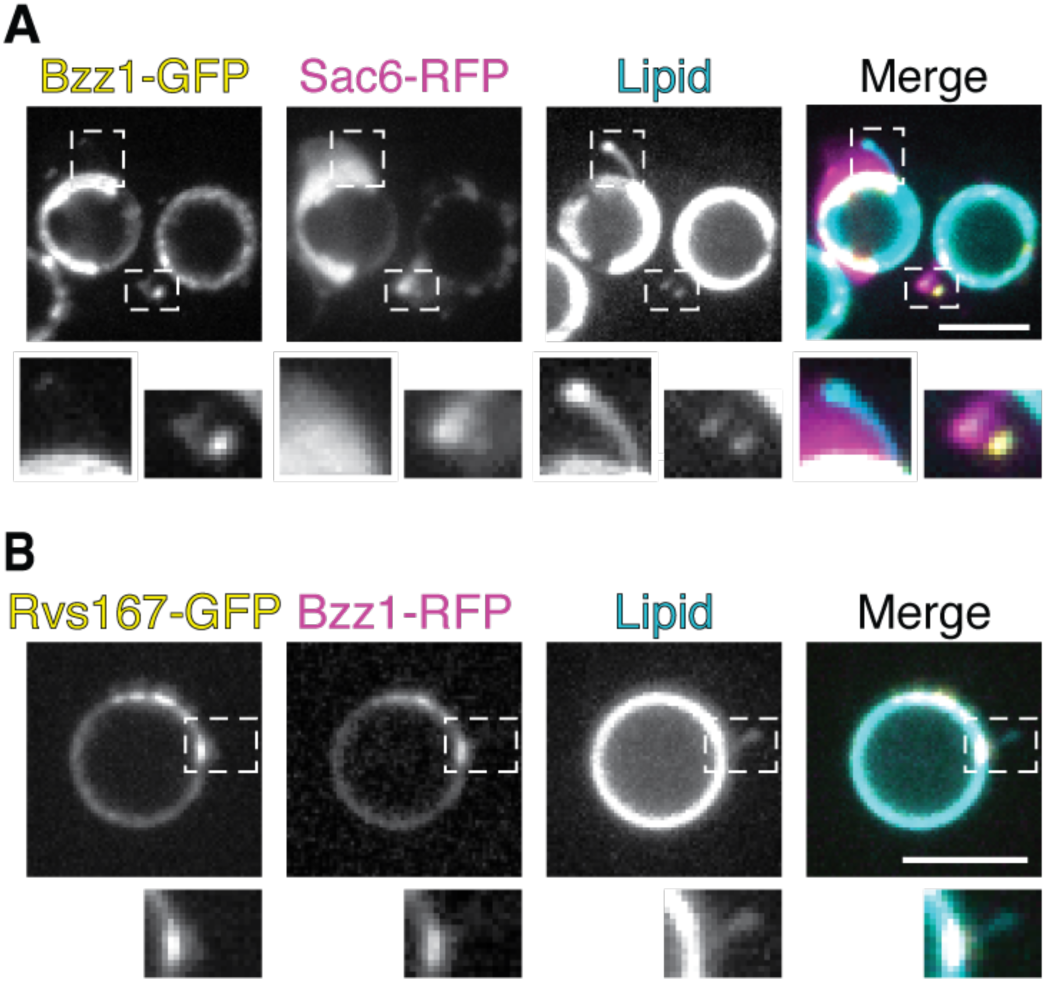
(A and B) scWASP-coated bilayers were incubated in cytoplasmic extract from strains expressing the indicated GFP- and RFP-tagged proteins. Maximum intensity projections of confocal stacks through the midsection of scWASP-coated bilayers containing lipid dye (0.5% MarinaBlue-DHPE) are displayed. Scale bars, 5 µm. (A) Insets show regions of an actin tail where membrane tubulation and membrane deformation occurred (lipid channel). (B) Inset shows membrane tubulation (lipid channel) at plaques of Rvs167-GFP and Bzz1-RFP accumulation.

**Figure S4.**
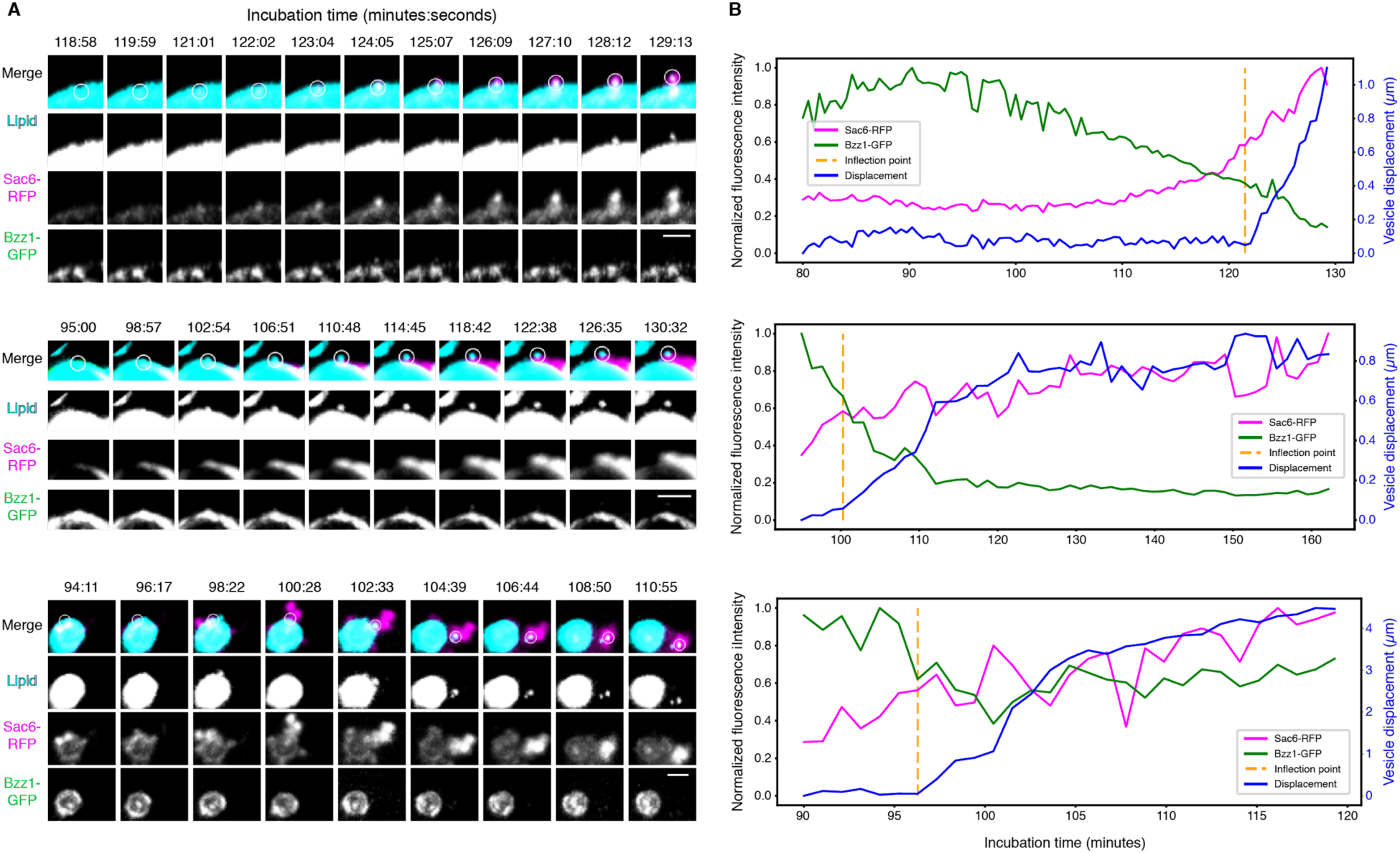
scWASP-coated supported lipid bilayers (with 2% Atto647-DOPE) were incubated in Bzz1-GFP Sac6-RFP cytoplasmic extract for the indicated times. (A) Montages of vesiculation events on 5 µm (upper two panels) and 2 µm (lower panel) beads. A maximum intensity projection of a substack through the center of the supported lipid bilayer is displayed for all channels. The white circle in the merged channel indicates the region used for quantitative analysis. Scale bars, 2 µm. (B) Quantification traces of fluorescence intensities and vesicle displacement from the vesiculation events visualized in (A).

**Figure S5.**
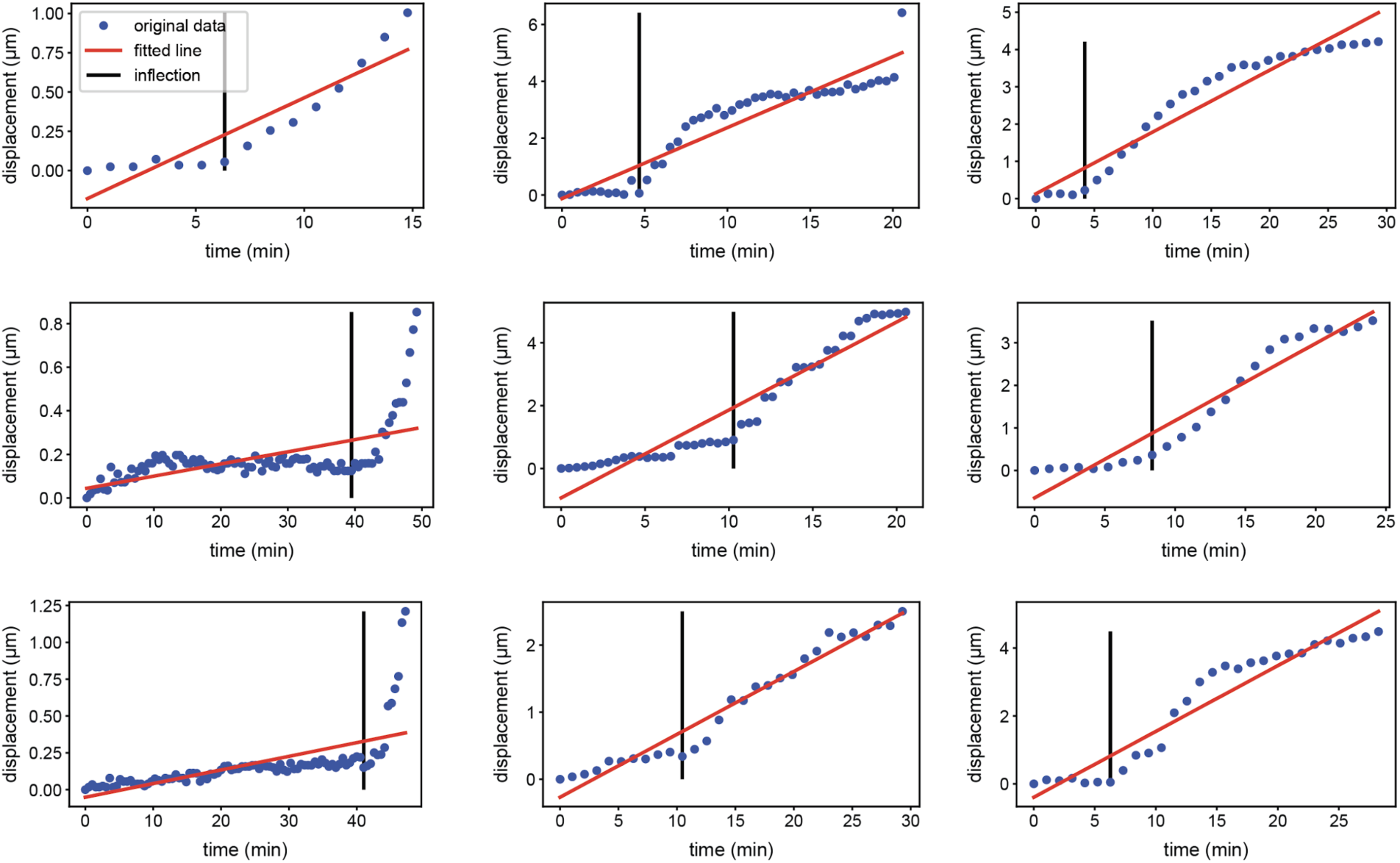
Gallery of randomly selected individual vesiculation event traces used for inflection point calculations. Blue dots represent the vesicle displacement in 3D volume from the origin position at the beginning of the event. A linear regression was fit to each vesiculation event trace. The inflection point was then determined as the time point at which the vesicle displacement was the maximum negative difference from the linear regression, out of all the time points before the point at the maximum positive difference from the linear regression. The constraint for points before the maximum positive difference from the linear regression was necessary to filter out points in displacement traces that plateaued at later time points.

**Figure S6.**
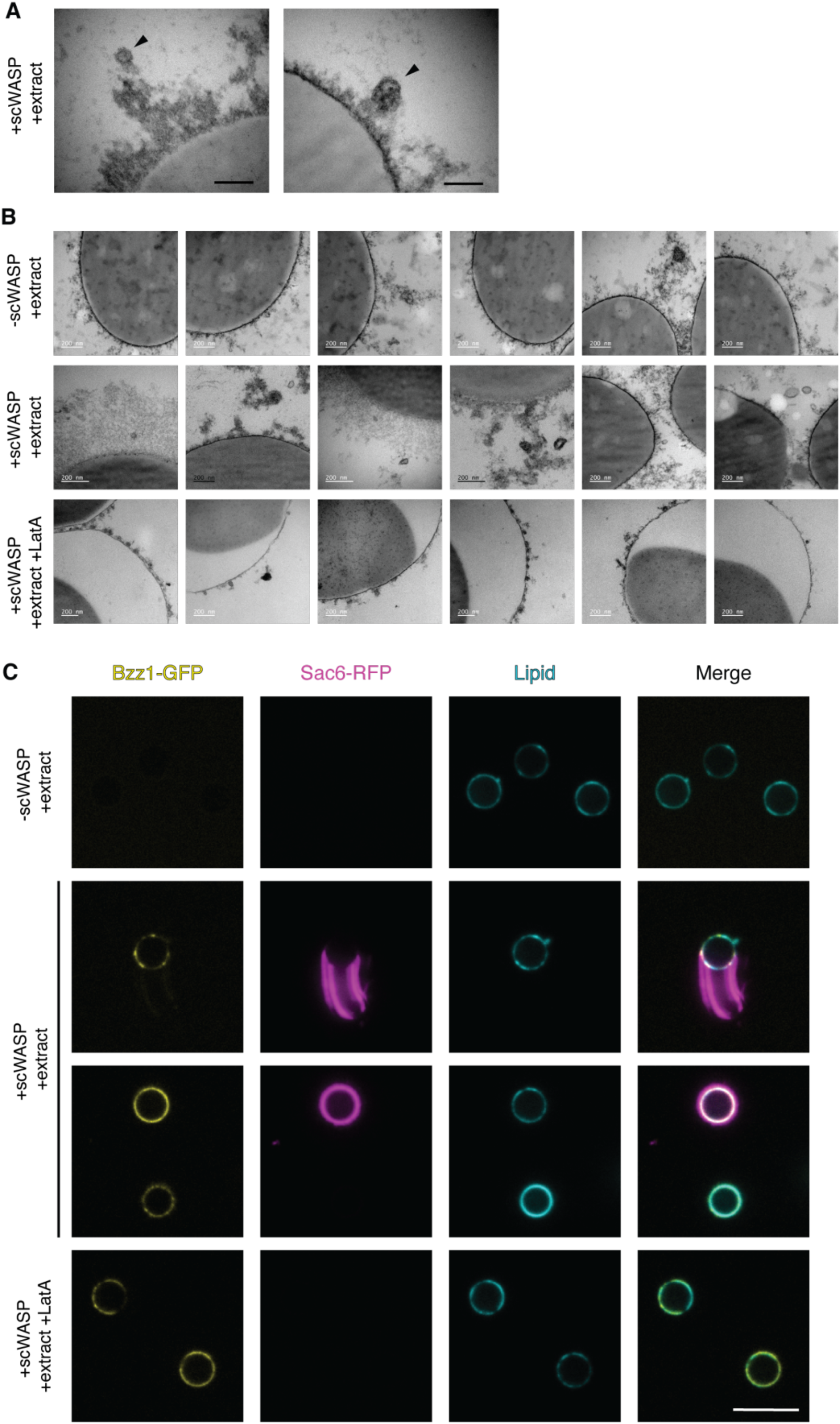
(A) High magnification EM images of vesicles (arrowheads) observed around supported lipid bilayers coated with scWASP and incubated in cytoplasmic extract. Scale bar, 100 nm. (B) Supported lipid bilayers on microbeads were either coated with or not coated with scWASP and incubated in cytoplasmic extract with or without latrunculin A, as indicated. Representative EM images of membrane deformations observed for each condition are presented here. (C) scWASP-coated bilayers were incubated in cytoplasmic extract from a strain expressing Bzz1-GFP and Sac6-RFP. Single confocal slices at the midsection of scWASP-coated bilayers containing lipid dye (2% Atto647-DOPE) are displayed. Scale bar, 10 µm.

**Table S1.**
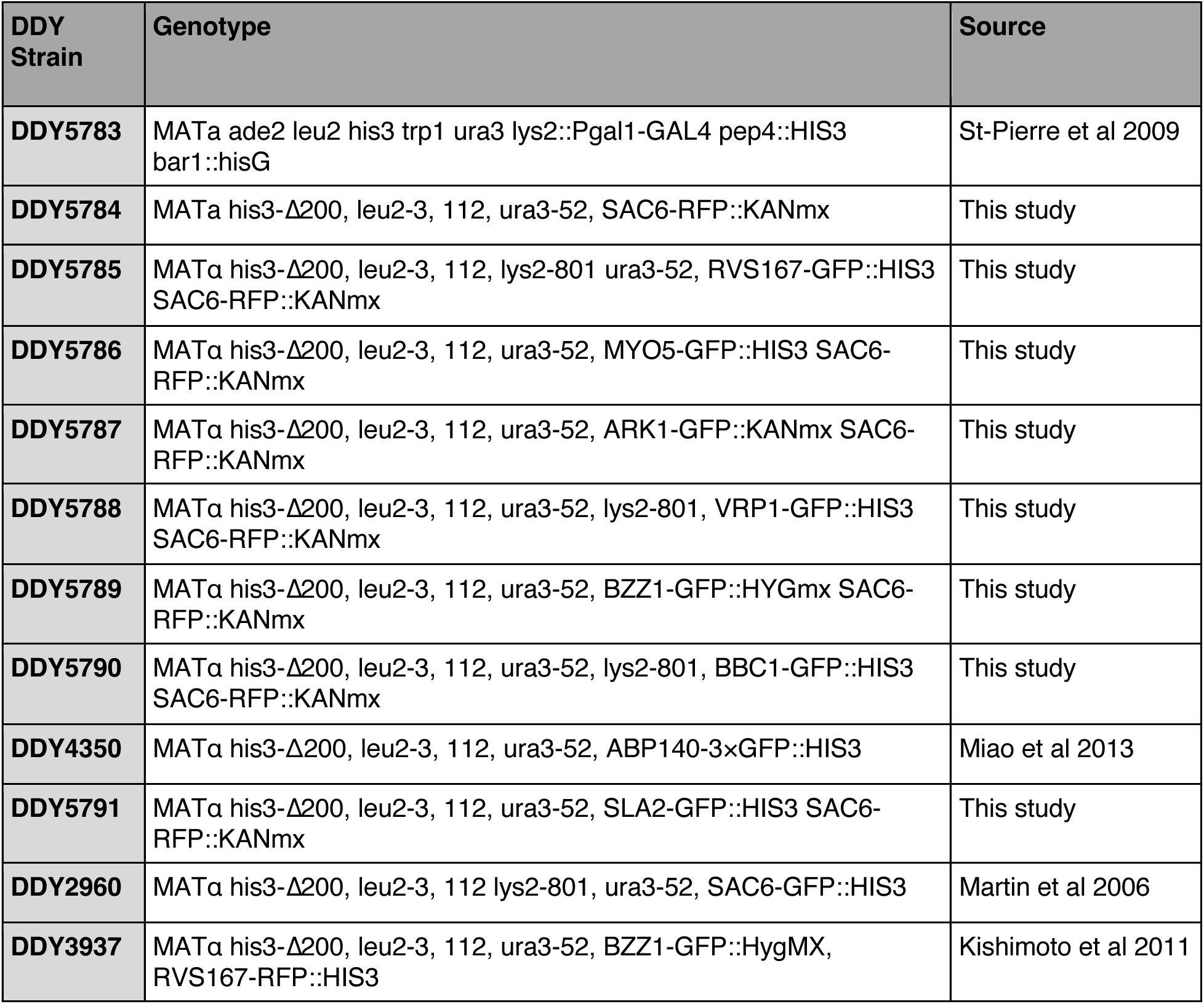
Strains used in this study.

**Table S2.** Manual quantification of the occurrence of actin-associated lipid protrusions on bilayers of differing lipid compositions. Where indicated, 20% phosphatidylserine (PS) and/or 5% phosphatidylinositol 4,5 bisphosphate (PI(4,5)P_2_) were added to bilayers containing 5% DGS-NTA, 2% ATTO 647-DOPE, and phosphatidylcholine (PC) up to 100%.

| **Lipid composition** | **n beads** | **Beads with actin-associated vesiculation** | **Percent** |
| --- | --- | --- | --- |
| **PC** | 103 | 3 | 0.029 |
| **PC + PS** | 154 | 11 | 0.071 |
| **PC + PI(4,5)P<sub>2</sub>** | 100 | 20 | 0.200 |
| **PC + PS + PI(4,5)P<sub>2</sub>** | 115 | 22 | 0.191 |

**Table S3.** Quantification of vesicle budding events observed in time lapse imaging. For each condition listed, the number of fluorescent vesicles formed from the surface of bead SLBs (budding events) was manually counted over the course of 30- to 60-minute 3D time-lapse experiments. The number of budding events per experiment was divided by the number of beads and averaged among replicate experiments to calculate the mean and standard deviation budding events per bead.

| Condition | n beads | Mean budding events per bead | Standard deviation |
| --- | --- | --- | --- |
| -scWASP -LatA +extract | 49 | 0.0476 | 0.0825 |
| +scWASP -LatA +extract | 97 | 0.248 | 0.291 |
| +scWASP +LatA +extract | 188 | 0.0481 | 0.0638 |
| +scWASP -LatA -extract | 16 | 0 | 0 |

**Table S4.** Quantification of vesicles observed by EM. For each condition listed, the number of vesicles and membrane structures at given distances away from the bead surface was manually counted using 2D electron micrographs. Events were categorized as ‘adjacent’ if within 100 nm of the bead surface or ‘nonadjacent’ if between 100 nm and 1000 nm from the bead surface. Vesicles found greater than 1000 nm from a particular bead were not counted.

| Condition | n beads | Adjacent events (lipid bilayers 10 - 100 nm from bead surface) | Adjacent events per bead | Nonadjacent events (lipid bilayers 100 - 1000 nm from bead surface) | Nonadjacent events per bead |
| --- | --- | --- | --- | --- | --- |
| -scWASP -LatA +extract | 47 | 3 | 0.0638 | 2 | 0.0426 |
| +scWASP -LatA +extract | 51 | 11 | 0.216 | 32 | 0.627 |
| +scWASP +LatA +extract | 43 | 36 | 0.837 | 7 | 0.163 |
| +scWASP -LatA -extract | 16 | 0 | 0 | 0 | 0 |

